# Reciprocal adipose-heart regulation of NRAC

**DOI:** 10.64898/2026.01.26.701858

**Authors:** Zhiyao Fu, Chao Zhang, Ren Zhang

**Affiliations:** Center for Molecular Medicine and Genetics, Wayne State University School of Medicine, Detroit, MI 48201, USA; Biostatistics Shared Resource, Winship Cancer Institute of Emory University, 718 Gatewood Rd. NE, Atlanta, GA 30322, USA

**Keywords:** adipocyte, adipose, CD36, fatty acids, heart, myocyte, NRAC

## Abstract

Fatty acids (FAs) are a major energy substrate, and their storage and utilization must be balanced across tissues to maintain metabolic homeostasis. White adipose tissue (WAT) and the heart play opposing roles in FA metabolism, with WAT serving as the primary site of FA storage and the heart relying heavily on FAs as an energy source. However, the molecular factors regulating FA handling between the two tissues remain incompletely understood. We previously identified NRAC as a nutritionally regulated adipose- and cardiac-enriched protein; however, how its expression is regulated between these tissues across nutritional states, and how this relates to systemic FA homeostasis, remains unclear. Here, we integrate analyses of human transcriptomic datasets with nutritional and genetic perturbations in mice to examine the association between NRAC and FA metabolism. Human single-cell transcriptomic data from GTEx show enrichment of NRAC expression in adipocytes and cardiomyocytes, and analysis of subcutaneous adipose tissue from the UK Twins cohort reveals strong associations between NRAC expression and the abundance of multiple adipose lipid species. In mice, NRAC expression in WAT and heart was reciprocally regulated by fasting and refeeding and showed opposing associations with circulating free fatty acid (FFA) levels; moreover, whole-body deletion of NRAC elevated circulating FFA. Together, these findings support a role for NRAC in nutritionally responsive inter-organ regulation of FA between WAT and the heart, with implications for systemic lipid homeostasis.

## Introduction

Metabolic diseases such as obesity and related cardiometabolic disorders are characterized by profound disturbances in lipid metabolism and fatty acid handling [1]. Dysregulated fatty acid mobilization, transport, and utilization contribute to insulin resistance, ectopic lipid accumulation, and impaired metabolic flexibility [2,3]. Understanding how fatty acid flux is coordinated between tissues in response to nutritional state remains a central challenge in metabolic research [4].

White adipose tissue (WAT) and the heart play central yet opposing roles in systemic fatty acid metabolism. During fasting, adipose tissue mobilizes stored triglycerides and releases fatty acids into the circulation, whereas the heart increases fatty acid uptake and oxidation to sustain its continuous energy demand [5]. In contrast, during feeding, adipose tissue promotes lipid storage while cardiac reliance on fatty acids is reduced [6]. Lipoprotein lipase (LPL) is a rate-limiting enzyme that hydrolyzes circulating triglyceride-rich lipoproteins into free fatty acids, and tissue-specific regulation of LPL activity is critical for partitioning triglyceride-derived fatty acids between storage tissues such as adipose and oxidative tissues such as the heart [7,8]. However, following LPL–mediated hydrolysis, the mechanisms governing fatty acid uptake and trafficking into target tissues, and how these processes are coordinately regulated across nutritional states, remain incompletely defined.

We previously identified NRAC (nutritionally regulated adipose and cardiac enriched protein) as a gene whose expression is highly enriched in adipose tissue and heart in mice [9]. This expression pattern is intriguing given the opposing roles of these tissues in fatty acid handling during fed and fasted states and raises the possibility that NRAC may participate in coordinating lipid flux between lipid storage and lipid utilization organs. Recent studies have implicated NRAC in fatty acid transport, and its genetic variations have been associated with obesity-related traits in humans, supporting its relevance to systemic lipid metabolism [10].

Despite these advances, important questions remain unanswered. How NRAC expression is coordinately regulated between adipose tissue and heart, two tissues with opposing metabolic roles, in response to nutritional state, how NRAC expression in these tissues relates to circulating FFA levels, and whether NRAC contributes to the regulation of systemic fatty acid homeostasis have not been clearly defined. Addressing these gaps is necessary to better understand how fatty acid transport is dynamically matched to tissue-specific metabolic demands. In the present study, we sought to define the tissue- and cell-type specificity of NRAC expression, examine its regulation by fasting and feeding, and assess its relationship to systemic fatty acid homeostasis using complementary human expression analyses and mouse genetic models. We provide the first evidence that NRAC is reciprocally regulated between WAT and heart by nutritional state, and that this regulation is linked to circulating fatty acid levels in vivo.

## Materials and Methods

### Mice

Mice were housed in a temperature-controlled facility (22–24 °C) under a 14-hour light/10-hour dark cycle and were provided ad libitum access to water and a standard chow diet (6% of calories from fat; Teklad 8664, Harlan Teklad, Indianapolis, IN), unless otherwise indicated. For fasting experiments, 8-week-old mice were subjected to a 24-hour fast, with age-matched fed mice maintained on ad libitum chow as controls. For refeeding studies, mice were fasted for 24 hours and subsequently refed chow diet for the indicated durations. NRAC knockout (KO) mice carrying the knockout-first allele 530016L24Rik^tm259628(L1L2_Bact_P)^ were obtained from the International Mouse Phenotyping Consortium (IMPC) [11]. For serum non-esterified fatty acid (NEFA) measurements, overnight-fasted NRAC KO mice and WT littermate controls were analyzed. Both male and female mice were included, as indicated in the figure legends. All animal protocols were approved by the Animal Care and Use Committee of Wayne State University.

### RNA extraction, quantitative real-time PCR, and measurement of free fatty acids

Dissected tissues were immediately placed into RNAlater solution (Ambion, Austin, TX) for subsequent RNA extraction. Total RNA was isolated from tissues with RNeasy Tissue Mini Kit with deoxyribonuclease treatment (Qiagen, Valencia, CA). One microgram of RNA was reverse transcribed to cDNA using random hexamers (Superscript; Ambion). Relative expression levels were calculated, and β-actin was used as an internal control. Primer sequences for mouse Nrac were: forward, 5′-AGTCTCTCGCTCTAATTCCCAC-3′; reverse, 5′-ACTTCCTGTTACCATCCCTCTC-3′. Primer sequences for mouse β-actin were: forward, 5′-GTGACGTTGACATCCGTAAAGA-3′; reverse, 5′-GCCGGACTCATCGTACTCC-3. Serum NEFA concentrations were measured using a commercial Free Fatty Acid Assay Kit (Sigma-Aldrich), according to the manufacturer’s instructions. All methods were carried out in accordance with relevant guidelines and regulations.

### Statistical analysis

Data are presented as mean ± s.e.m. and statistical significance was assessed using unpaired two-tailed Student’s *t*-tests unless otherwise specified. For multi-group comparisons (fasting/refeeding), differences among groups were assessed by one-way ANOVA followed, when appropriate, by Fisher’s LSD post hoc test. Linear correlations were evaluated using Pearson’s correlation analysis. RNA-sequencing data were obtained from GTEx (v10) [12] to analyze NRAC expression across human tissues using bulk and single-cell datasets. Boxplots display the interquartile range (IQR), with the horizontal line indicating the median; whiskers extend to 1.5 × IQR, and outlier samples are shown as individual data points. All statistical analyses were performed using SAS version 9.4 (SAS Institute, Cary, NC, USA). Figures were generated using JASP. Differences were considered statistically significant at P < 0.05.

## Results

### NRAC expression is highly enriched in human adipose and heart tissues and is specific to adipocytes and myocytes

In 2012, we reported that NRAC expression is highly enriched in adipose tissue and heart across mouse tissues, at a time when Genotype-Tissue Expression (GTEx) data were not yet available. The GTEx project now enables a comprehensive and unbiased evaluation of gene expression across human tissues [12]. Examining NRAC expression in GTEx is important for several reasons: (i) to assess cross-species conservation of tissue specificity, and (ii) to leverage single-cell transcriptomic data to resolve cell-type–specific expression within complex tissues.

To define the tissue distribution of human NRAC, we analyzed bulk RNA-seq data from GTEx version 10. Across 54 human tissues, NRAC expression was highly enriched in adipose and cardiovascular tissues (Fig. 1A). Among adipose depots, NRAC showed robust expression in subcutaneous and visceral adipose tissues, and breast mammary tissue. In the cardiovascular system, high NRAC expression was observed in the heart atrial appendage, left ventricle, and coronary artery. In contrast, NRAC expression was low or nearly undetectable in the majority of other tissues, indicating a highly tissue-restricted expression pattern in humans.

**Figure 1.**
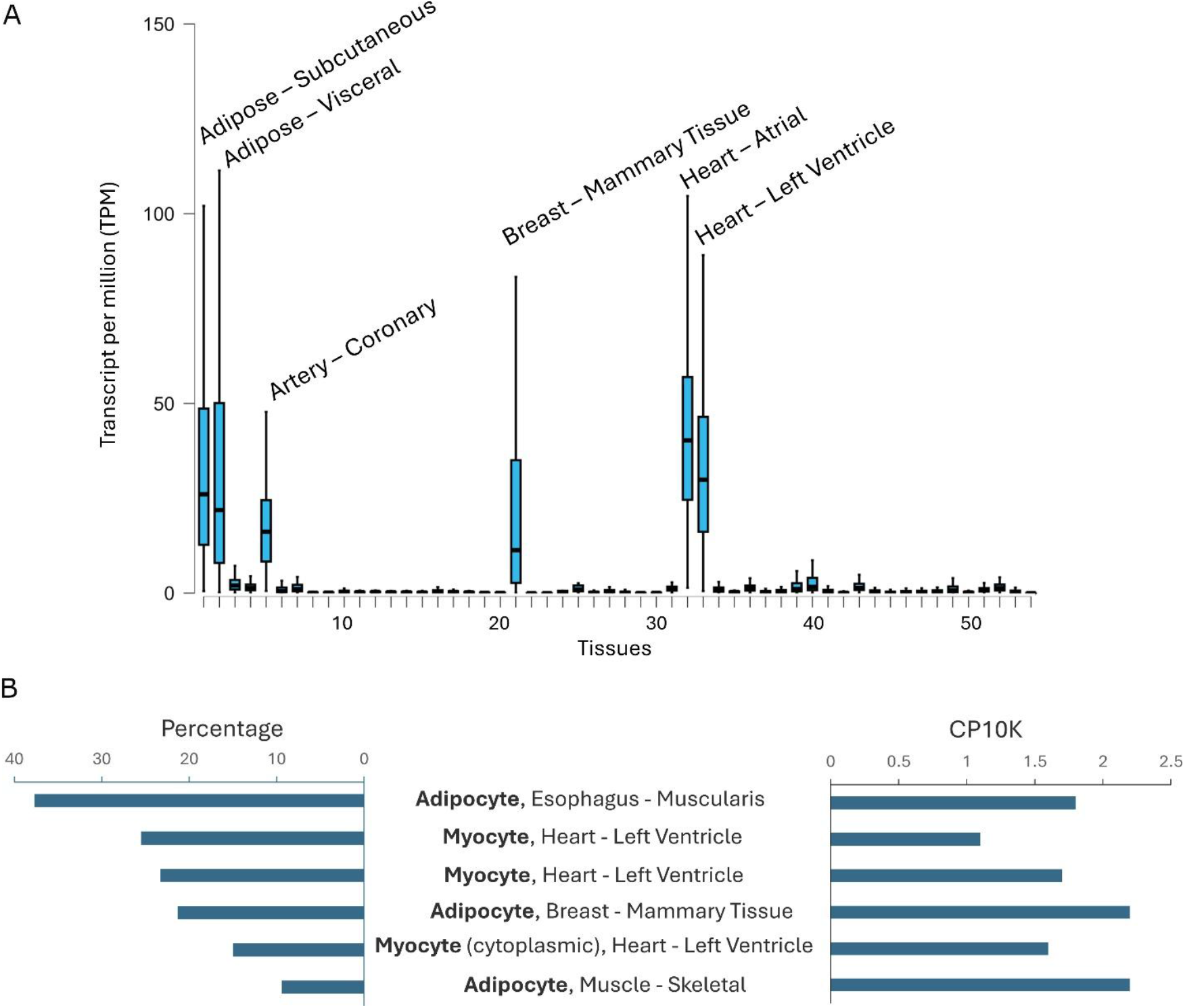
Human NRAC expression is specific to adipocytes and myocytes in adipose and heart tissues. (A) NRAC expression is highly enriched in human adipose and heart tissues. Boxplots show bulk RNA-seq expression of NRAC (TPM) across 54 human tissues from the GTEx v10 dataset. Each point represents an individual donor sample. The boxes indicate the interquartile range (IQR), the horizontal line denotes the median, and whiskers extend to 1.5×IQR; outlier samples are shown as individual points. NRAC expression is highly enriched in adipose tissue and heart, with minimal expression in most other tissues. TPM = transcripts per million. The tissue indices correspond to the following tissues (in plotted order): 1) Adipose – Subcutaneous; 2) Adipose – Visceral; 3) Adrenal Gland; 4) Artery – Aorta; 5) Artery Coronary; 6) Artery – Tibial; 7) Bladder; 8) Brain – Amygdala; 9) Brain – Anterior Cingulate Cortex (BA24); 10) Brain – Caudate (Basal Ganglia); 11) Brain – Cerebellar Hemisphere; 12) Brain – Cerebellum; 13) Brain – Cortex; 14) Brain – Frontal Cortex; 15) Brain – Hippocampus; 16) Brain – Hypothalamus; 17) Brain – Nucleus Accumbens; 18) Brain – Putamen; 19) Brain – Spinal Cord; 20) Brain – Substantia Nigra; 21) Breast – Mammary Tissue; 22) Cells – Cultured Fibroblasts; 23) Cells – EBV-Transformed Lymphocytes; 24) Cervix – Ectocervix; 25) Cervix – Endocervix; 26) Colon – Sigmoid; 27) Colon – Transverse; 28) Esophagus – Gastroesophageal Junction; 29) Esophagus – Mucosa; 30) Esophagus – Muscularis; 31) Fallopian Tube; 32) Heart Atrial Appendage; 33) Heart – Left Ventricle; 34) Kidney – Cortex; 35) Kidney – Medulla; 36) Liver; 37) Lung; 38) Minor Salivary Gland; 39) Muscle – Skeletal; 40) Nerve – Tibial; 41) Ovary; 42) Pancreas; 43) Pituitary; 44) Prostate; 45) Skin – Not Sun-Exposed; 46) Skin – Sun-Exposed; 47) Small Intestine – Terminal Ileum; 48) Spleen; 49) Stomach; 50) Testis; 51) Thyroid; 52) Uterus; 53) Vagina; 54) Whole Blood. (B) Single-cell RNA-seq analysis reveals NRAC expression is specific to adipocytes and cardiomyocytes. Bar plots summarize the six cell types that rank highest for both the percentage of NRAC-expressing cells (prevalence) and expression intensity (CP10K) based on GTEx v10 single-cell RNA-sequencing datasets. The left panel shows the percentage of cells expressing NRAC within each indicated cell type, and the right panel shows the average expression level among expressing cells. NRAC expression is specific in adipocytes and myocytes, showing pronounced cell-type specificity within adipose and heart tissues. For CP10K-based ranking, cell types with fewer than 1% of NRAC-expressing cells were excluded to reduce sampling noise. CP10K = counts per 10,000.

Because bulk tissue expression reflects the aggregate signal from multiple cell types, we next examined single-cell RNA-seq data from GTEx to determine the cellular specificity of NRAC expression. Adipose and heart tissues contain diverse cell populations, including immune cells, endothelial cells, fibroblasts, and stromal cells in addition to adipocytes and myocytes. We therefore evaluated NRAC expression using two commonly used single-cell metrics: the percentage of cells expressing NRAC and normalized expression levels (counts per 10,000 reads; CP10K). Across all annotated cell types, NRAC expression was highly restricted to adipocytes and myocytes (Fig.1B). Ranking cell types by either the percentage of NRAC-positive cells or CP10K yielded the same top six cell populations, all of which corresponded to adipocytes or myocytes.

Together, these data demonstrate that NRAC expression in humans is highly enriched in adipose and heart tissues and is specifically localized to adipocytes and myocytes at the cellular level. This conserved tissue and cell-type specificity between mice and humans supports a fundamental role for NRAC in adipocyte and myocyte biology.

### Reciprocal regulation of NRAC expression in white adipose tissue and heart

WAT and heart play complementary and often reciprocal roles in lipid utilization in response to nutritional state. During fasting, adipose tissue mobilizes stored triglycerides, whereas the heart increases reliance on fatty acid uptake and oxidation to meet its energetic demands. Based on the adipocyte- and myocyte-specific expression of NRAC, we hypothesized that NRAC expression is reciprocally regulated in WAT and heart in response to fasting and refeeding. To test this hypothesis, we examined NRAC expression following a 24-h fast and during the refeeding period.

In WAT, a 24-h fast significantly reduced NRAC expression compared with the fed state (Fig. 2A). Refeeding led to a rapid and progressive restoration of NRAC expression, with significant increases observed at 3 h and 6 h. These results indicate that NRAC expression in adipose tissue is nutrient-responsive and suppressed during prolonged fasting. In contrast, NRAC expression in the heart was reciprocally regulated (Fig. 2B). A 24-h fast markedly increased cardiac NRAC expression relative to the fed state. Upon refeeding, NRAC expression declined in a time-dependent manner, with significant reductions evident as early as 1 h and continuing through 3 h and 6 h of refeeding, approaching or falling below fed levels. To further evaluate the reciprocal relationship between adipose and cardiac NRAC expression, we performed correlation analysis across individual animals. NRAC expression in WAT and heart exhibited a strong inverse correlation (r = −0.69, P < 0.001) (Fig. 2C), providing quantitative support for reciprocal regulation across nutritional states.

**Fig. 2.**
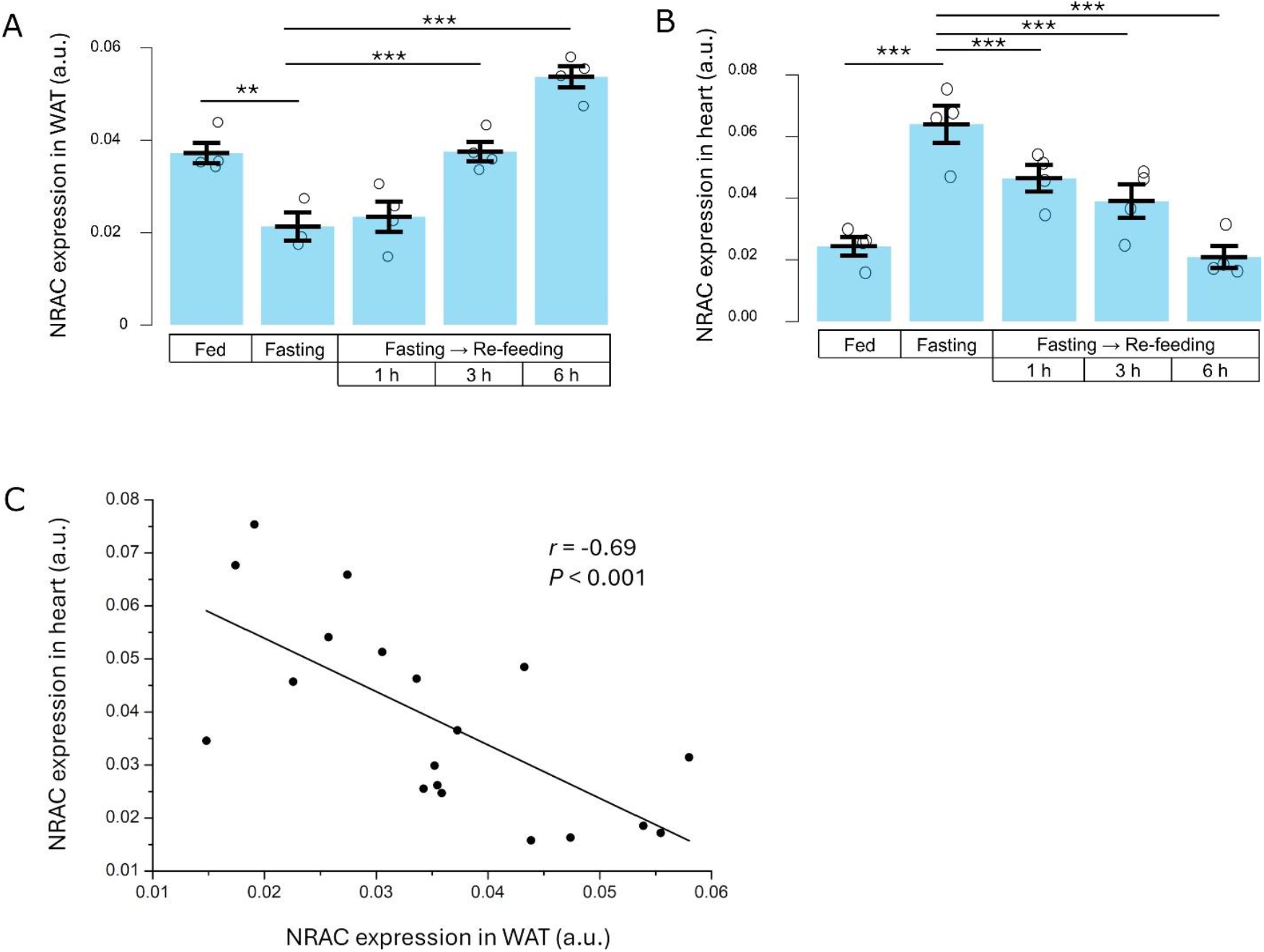
Reciprocal regulation of NRAC expression in white adipose tissue and heart. (A) NRAC expression in white adipose tissue (WAT) and (B) heart from mice under fed, 24-h fasted, and re-fed conditions (1, 3, and 6 h after re-feeding). Bars represent mean ± SEM, and each open circle represents one individual mouse. (C) Correlation between NRAC expression in WAT and heart across all nutritional conditions. Each dot represents one mouse. Solid line indicates linear regression. Correlation was assessed by Pearson correlation (r = -0.69, P < 0.001). Statistical significance in (A) and (B) was determined by one-way ANOVA (*F*=24.7; *P*<0.001 and *F*=25.2; *P*<0.001 for A and B, respectively) with LSD *post hoc* multiple-comparison testing. N = 4 per group. P < 0.01 (**), P < 0.001 (***).

Together, these findings demonstrate that NRAC expression is reciprocally regulated in WAT and heart in response to fasting and refeeding, mirroring the opposing metabolic roles of these tissues in lipid mobilization and utilization. This dynamic regulation suggests that NRAC may function as a coordinated metabolic regulator linking adipose lipid release with cardiac fatty acid demand.

### Opposing associations of NRAC expression in WAT and heart with circulating fatty acids

Adipose tissue and heart exhibit opposing roles in systemic fatty acid handling, with adipose tissue serving as the primary site of fatty acid release and the heart acting as a major site of fatty acid uptake and utilization. Recent studies have identified NRAC as a critical regulator of fatty acid transport, suggesting that its expression may be linked to circulating fatty acid availability. We therefore hypothesized that NRAC expression in adipose tissue and heart would be differentially associated with serum non-esterified fatty acid (NEFA) levels. To test this hypothesis, we examined the relationship between NRAC expression and serum NEFA concentrations in mice subjected to fasting and refeeding.

NRAC expression in WAT was inversely correlated with circulating NEFA levels (r = −0.49, P < 0.05) (Fig. 3A). Animals with higher NRAC expression in WAT exhibited lower serum NEFA concentrations, whereas reduced NRAC expression was associated with elevated NEFA levels. This relationship is consistent with a role for NRAC in adipose tissue that restrains fatty acid release or promotes intracellular fatty acid handling under nutrient-replete conditions. In contrast, NRAC expression in the heart showed a positive association with serum NEFA levels (r = 0.52, P < 0.05) (Fig. 3B). Higher cardiac NRAC expression correlated with increased circulating NEFA concentrations, consistent with enhanced fatty acid uptake and utilization by the heart during states of elevated energy demand.

**Fig. 3.**
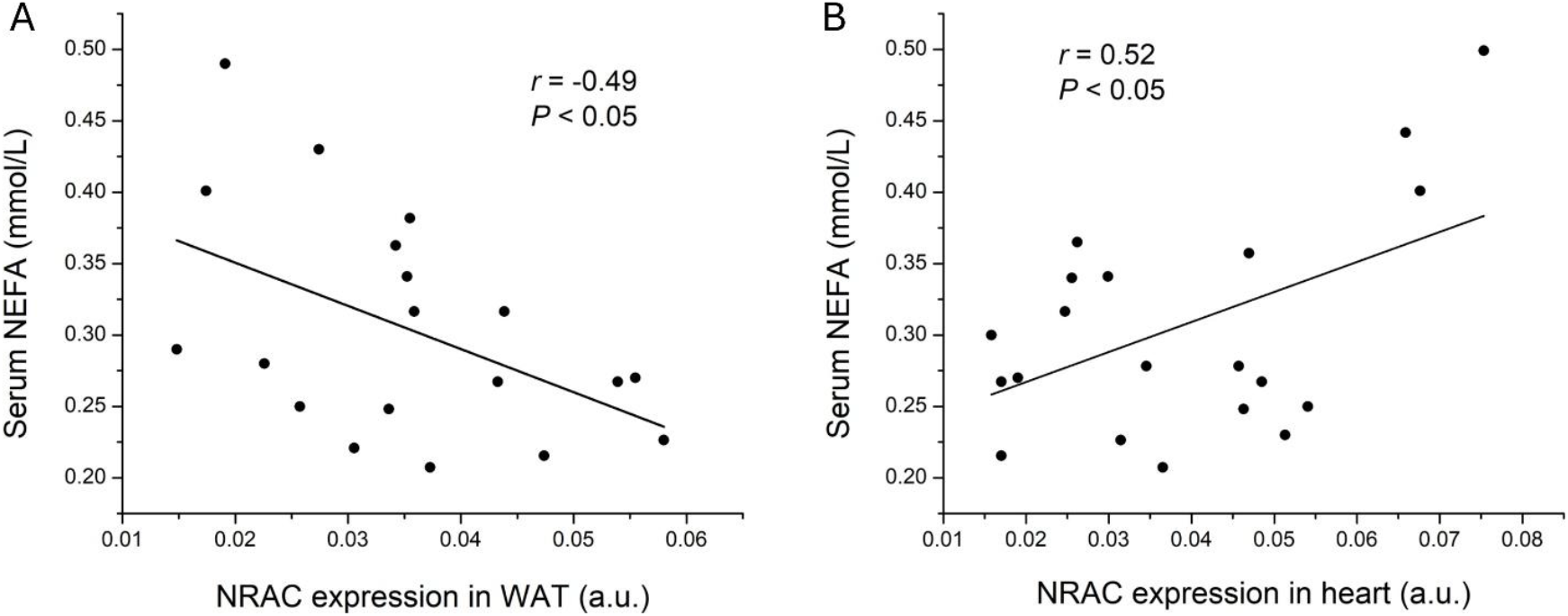
Opposing associations of NRAC expression in WAT and heart with circulating fatty acids. A) Correlation between NRAC expression in white adipose tissue (WAT) and serum non-esterified fatty acid (NEFA) concentrations. B) Correlation between NRAC expression in heart and serum NEFA concentrations. Each dot represents one individual mouse. Solid lines indicate linear regression. Correlations were assessed using Pearson correlation analysis, with correlation coefficients (r) and P values indicated on the plots.

Together, these findings reveal opposing associations between NRAC expression in adipose tissue and heart with circulating fatty acids, reinforcing the concept of reciprocal NRAC regulation across tissues. These data support a model in which NRAC coordinates systemic lipid flux by coupling adipose fatty acid release with cardiac fatty acid utilization in response to metabolic state.

### NRAC deficiency elevates circulating non-esterified fatty acids in mice

To establish a causal role for NRAC in systemic fatty acid regulation, we examined circulating NEFA levels in NRAC knockout (KO) mice. Loss-of-function mouse models were generated by the International Mouse Phenotyping Consortium (IMPC) [11], and we utilized the Nrac knockout first allele A530016L24Rik^tm259628(L1L2_Bact_P)^. This allele results in global deletion of NRAC, enabling assessment of its role in whole-body lipid homeostasis. We hypothesized that NRAC deficiency would impair fatty acid uptake into tissues, resulting in elevated circulating NEFA levels. To test this hypothesis, serum NEFA concentrations were measured in overnight-fasted NRAC KO mice and WT littermate controls. Consistent with this hypothesis, NRAC deficiency significantly increased circulating NEFA levels in the KO mice. In male mice, NRAC KO animals exhibited a marked elevation in serum NEFA compared with WT controls (Fig. 4A). A similar increase in circulating NEFA was observed in female NRAC KO mice (Fig. 4B), indicating that the effect of NRAC loss on fatty acid homeostasis is not sex-dependent. Together, these results demonstrate that NRAC is needed for maintaining normal circulating fatty acid levels in vivo. The elevation of serum NEFA in NRAC-deficient mice supports a causal role for NRAC in regulating systemic fatty acid handling.

**Fig. 4.**
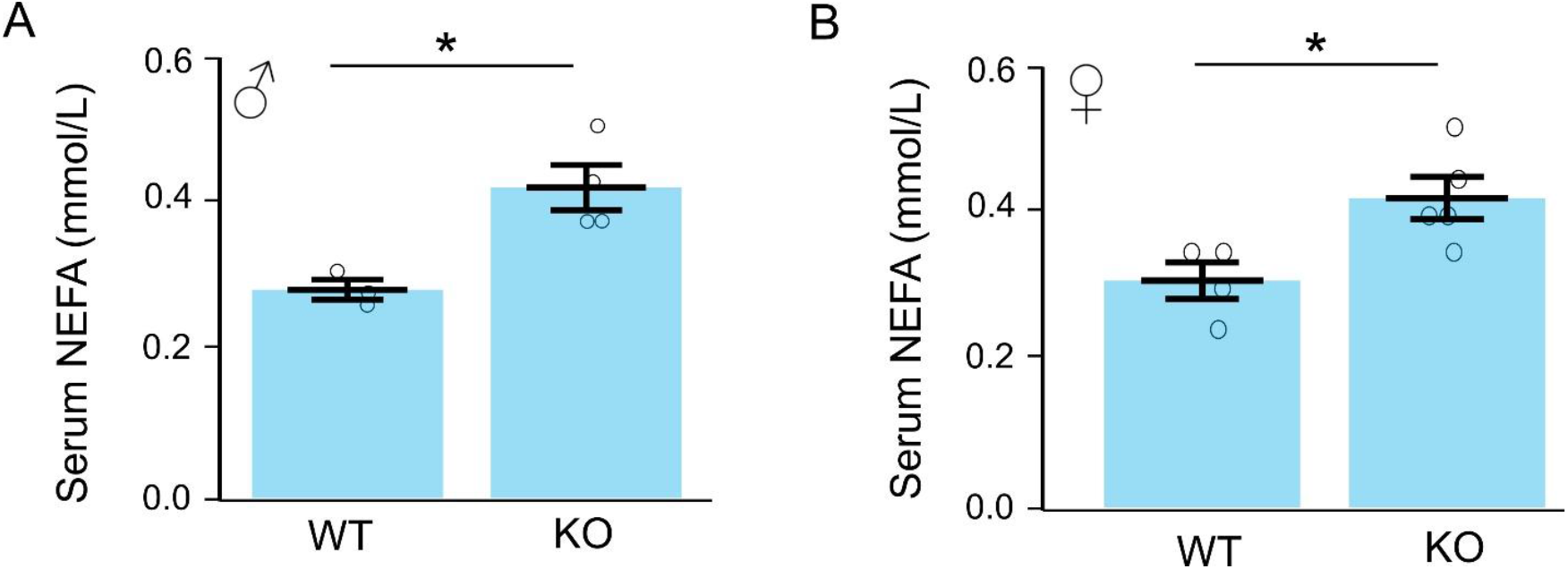
NRAC deficiency elevates circulating NEFA in mice. Serum non-esterified fatty acid (NEFA) concentrations in (A) male and (B) female wild-type (WT) and NRAC knockout (KO) mice. Data are represented as mean ± SEM. Dots represent individual animals. N = 4 - 5 per group. *, P < 0.05.

### Associations between adipose NRAC expression and cardiometabolic traits in a human cohort

To evaluate the relevance of NRAC to human cardiometabolic phenotypes, we examined associations between adipose NRAC expression (ENSG00000184601) and DXA-derived fat-distribution traits, circulating lipids, and cardiac electrical measures in the TwinsUK cohort [13]. Summary statistics were obtained from a transcriptome-wide association analysis of adipose RNA-sequencing data and cardiometabolic phenotypes.

Adipose NRAC expression showed exceptionally strong and coherent associations with central and visceral adiposity (Table 1). NRAC expression was inversely associated with the cross-sectional area of visceral fat within the abdominal cavity (β = −0.459, P = 1.67 × 10^−20^), the percentage of fat in the largest visceral fat region (β = −0.343, P = 2.69 × 10^−15^), and the ratio of visceral to gynoid fat (β = −0.362, P = 1.85 × 10^−26^), indicating that higher NRAC expression is strongly linked to reduced accumulation of metabolically deleterious organ-surrounding fat. Consistent with this, NRAC expression was also inversely associated with multiple measures of abdominal and trunk adiposity, including fat tissue in the trunk region (β = −0.534, P = 4.08 × 10^−17^), percentage of fat in the android region (β = −0.359, P = 1.18 × 10^−15^), percentage of trunk tissue that is fat (β = −0.336, P = 1.72 × 10^−13^), and the trunk-to-limb fat ratio (β = −0.328, P = 3.30 × 10^−27^). In contrast, adipose NRAC expression was positively associated with gynoid and lower-body depots, including all tissue in the gynoid region (β = 0.147, P = 0.007) and right leg mass (β = 0.229, P = 3.7 × 10^−6^), consistent with preferential partitioning of lipid toward metabolically protective peripheral depots.

**Table 1.** Associations between adipose NRAC expression and fat distribution, lipoproteins, and cardiac traits in the TwinsUK cohort.

| Domain | Trait | $\beta$ | P value |
| --- | --- | --- | --- |
| Visceral adiposity | Cross sectional area of fat inside abdominal cavity | -0.459 | 1.67E-20 |
|  | Percentage of fat in largest visceral fat region / percentage of fat in gynoid region | -0.362 | 1.85E-26 |
|  | Percentage of fat in largest visceral fat region | -0.343 | 2.69E-15 |
| Abdominal<br>(android/trunk)<br>adiposity | Fat tissue in trunk region | -0.534 | 4.08E-17 |
|  | Trunk fat / limb fat | -0.328 | 3.3E-27 |
|  | Percentage of fat in android region | -0.359 | 1.18E-15 |
|  | Percentage of fat tissue to total tissue mass in trunk region | -0.336 | 1.72E-13 |
| Total adiposity | Percentage of total body mass that is fat | -0.140 | 0.003 |
| Gynoid tissue | All tissue in gynoid region | 0.147 | 0.007 |
|  | Mass in right leg region | 0.229 | 3.7E-06 |
| Lipids | Serum Apolipoprotein B | -0.068 | 0.022 |
|  | Serum triglycerides | -0.183 | 3.63E-09 |
|  | High-density lipoprotein | 0.150 | 1.8E-06 |
| Heart | ECG P axis | 0.152 | 0.002 |

These fat-distribution effects were accompanied by a favorable circulating lipid profile. Adipose NRAC expression was inversely associated with serum apolipoprotein B (ApoB) (β = −0.068, P = 0.022) and serum triglycerides (β = −0.183, P = 3.63 × 10^−9^), and positively associated with high-density lipoprotein (HDL) levels (β = 0.150, P = 1.8 × 10^−6^), indicating reduced atherogenic lipoprotein burden and improved lipid handling in individuals with higher NRAC expression.

Although NRAC expression was also modestly inversely associated with total body fat percentage (β = −0.140, P = 0.003), the magnitude of its effects on regional fat distribution and lipoproteins was substantially greater, suggesting that NRAC primarily influences fat partitioning rather than overall adiposity.

Notably, adipose NRAC expression was also associated with a cardiac electrical phenotype, showing a positive association with the ECG P-axis (β = 0.152, P = 0.002), a measure reflecting atrial depolarization and cardiac electrical remodeling. This link to a cardiac trait provides further evidence that adipose NRAC expression is coupled to systemic cardiometabolic physiology.

Together, these data suggest that adipose NRAC expression is associated with reduced visceral and abdominal adiposity, increased gynoid fat depots, a more favorable lipoprotein profile, and healthier cardiac electrical traits, supporting a link between NRAC and cardiometabolic phenotypes in humans.

## Discussion

The ongoing obesity epidemic and its associated cardiometabolic complications highlight the importance of understanding how fatty acid metabolism is coordinated across tissues. Dysregulated fatty acid mobilization and uptake contribute to elevated circulating free fatty acids, ectopic lipid accumulation, insulin resistance, and cardiovascular dysfunction [1,2,4,5]. Although individual pathways governing lipid storage and oxidation have been extensively studied, how fatty acid transport is dynamically coordinated between metabolically opposing tissues remains incompletely understood.

In this study, we identify NRAC as a tissue- and cell-type–specific factor that links adipose tissue and heart fatty acid metabolism in a nutritionally responsive manner. Building on our earlier observation that NRAC expression is highly enriched in adipose tissue and heart, we show that this enrichment is conserved in humans and localized to adipocytes and myocytes. Importantly, NRAC expression is reciprocally regulated between WAT and heart during fasting and refeeding, paralleling the opposing metabolic roles of these tissues. In addition, NRAC expression exhibits opposing associations with circulating FFA levels in WAT and heart, and whole-body deletion of NRAC leads to elevated circulating FFA, supporting a role for NRAC in systemic fatty acid homeostasis.

Recent studies demonstrating a physical and functional interaction between NRAC and CD36 provide important mechanistic context for understanding NRAC function in lipid metabolism [10]. CD36 is a major fatty acid translocase whose activity is regulated primarily through dynamic trafficking between membrane compartments rather than by total expression levels [14]. Accumulating evidence indicates that NRAC does not function as a fatty acid transporter itself, but instead influences CD36-dependent fatty acid uptake by modulating CD36 trafficking and membrane behavior [10]. Importantly, CD36 internalization and recycling have been reported to occur through both caveolae-associated and clathrin-mediated pathways, and these routes differ in kinetics, regulation, and functional outcomes [10,15]. How NRAC interfaces with these distinct endocytic mechanisms remains incompletely defined and is likely to depend on cell type, metabolic state, and differentiation status. Consistent with this complexity, our human analyses show that NRAC expression is positively associated with some adipose depots and negatively associated with others, indicating that NRAC is linked to divergent patterns of fat distribution. Together with prior work identifying C14orf180 (NRAC) as an epigenetically regulated factor involved in adipocyte differentiation and lipid storage [16], these findings support a view in which NRAC contributes to fatty-acid handling in a tissue-specific and context-dependent manner, potentially through differential engagement of caveolae- and clathrin-dependent CD36 trafficking pathways.

Our previous mouse studies showed that adipose NRAC expression is markedly reduced in obesity, both in diet-induced and genetically obese (ob/ob) models, establishing a robust negative relationship between NRAC and adiposity [9]. This inverse association is conserved in humans. In the TwinsUK cohort, adipose NRAC expression is strongly and inversely associated with multiple measures of visceral and abdominal fat, including visceral fat area, trunk fat mass, android fat percentage, and central fat ratios. Because circulating FFA levels are typically elevated in states of increased central adiposity, these human associations are compatible with a broader relationship between adipose NRAC expression and lipid-rich metabolic environments. In contrast, adipose NRAC expression shows positive associations with gynoid and lower-body depots, including hip–thigh tissue mass and leg mass. Gynoid fat is widely regarded as a metabolically protective fat compartment, in contrast to visceral fat, suggesting that NRAC is linked to differential fat partitioning rather than simply total fat mass. Consistent with this interpretation, higher adipose NRAC expression is also associated with a more favorable lipoprotein profile, including lower apolipoprotein B and triglycerides and higher HDL, as well as a cardiac electrical phenotype (ECG P-axis) associated with healthier atrial remodeling. Together, these human associations support a conserved relationship between adipose NRAC expression, fat distribution, and cardiometabolic traits, extending our mouse findings to a clinically relevant human context.

Several limitations of the present study should be acknowledged. Our analyses primarily focused on NRAC mRNA expression, and protein abundance, post-translational regulation, and subcellular localization were not directly assessed. In addition, while whole-body NRAC deletion establishes a role in systemic fatty acid homeostasis, tissue-specific genetic models will be required to define the relative contributions of adipose tissue and heart. Finally, direct measurements of fatty acid uptake at the tissue level will be necessary to further validate the proposed model.

These limitations highlight important directions for future research. Defining NRAC protein localization, its dynamic interaction with CD36, and its regulation by hormonal and nutritional cues will be critical to elucidating its molecular mechanism. Tissue-specific gain- and loss-of-function approaches will further clarify how NRAC integrates adipose–heart cross-talk. Given the conservation of NRAC expression in humans, exploring its relevance in human metabolic disease may also provide translational insight.

In summary, this study identifies NRAC as a nutritionally regulated factor expressed in adipose tissue and heart that is associated with tissue-specific fatty-acid handling and systemic lipid homeostasis. Our findings, together with prior mechanistic studies linking NRAC to CD36, suggest that NRAC may influence fatty-acid uptake capacity in a context-dependent manner, potentially through modulation of CD36 trafficking. Disruption of NRAC-dependent regulation may therefore contribute to lipid dysregulation in cardiometabolic disease.

## Acknowledgements

This work was supported in part by the National Institutes of Health grant DK132065 (to R.Z.).

